# CandyCollect: At-home saliva sampling for capture of respiratory pathogens

**DOI:** 10.1101/2021.10.19.464911

**Authors:** Ulri N. Lee, Xiaojing Su, Damielle L. Hieber, Wan-chen Tu, Anika M. McManamen, Meg G. Takezawa, Grant W. Hassan, Tung Ching Chan, Karen N. Adams, Ellen R. Wald, Gregory P. DeMuri, Erwin Berthier, Ashleigh B. Theberge, Sanitta Thongpang

## Abstract

*Streptococcus pyogenes* is a major human-specific bacterial pathogen and a common cause of a wide range of symptoms from mild infection such as pharyngitis (commonly called strep throat) to life-threatening invasive infection and post-infectious sequelae. Traditional methods for diagnosis include collecting a sample using a pharyngeal swab, which can cause discomfort and even discourage adults and children from seeking proper testing and treatment in the clinic. Saliva samples are an alternative to pharyngeal swabs. To improve the testing experience for strep throat, we developed a novel lollipop-inspired sampling platform (called CandyCollect) to capture bacteria in saliva. The device can be used in clinics or in the home and shipped back to a lab for analysis, integrating with telemedicine. CandyCollect is designed to capture bacteria on an oxygen plasma treated polystyrene surface embedded with flavoring substances to enhance the experience for children and inform the required time to complete the sampling process. In addition, the open channel structure prevents the tongue from scraping and removing the captured bacteria. Flavoring substances did not affect bacterial capture and the device has a shelf life of at least 2 months (with experiments ongoing to extend the shelf life). We performed a usability study with 17 participants who provided feedback on the device design and the dissolving time of the candy. This technology and advanced processing techniques, including polymerase chain reaction (PCR), will enable user-friendly and effective diagnosis of streptococcal pharyngitis.

## Introduction

Acute respiratory infections are common with children (average 3-5 upper respiratory infections per year), and may lead to additional complications if left untreated.^1^ In developing countries, these illnesses can be more severe, with gaps in healthcare increasing the risk of morbidity and mortality.^2^ Additionally, even in developed countries, people who have limited access to clinics, such as in rural areas, or those without insurance may also be unable to receive appropriate diagnosis and treatment. Here we develop a new sample collection platform for a common bacterial infection, group A streptococcal (GAS) pharyngitis, also known as strep throat, caused by *Streptococcus pyogenes*. Pharyngitis is one of the most common reasons for which children and young adults seek medical care and the Group A streptococcus (GAS) is the most frequent cause of bacterial pharyngitis in these age groups.^3^ GAS pharyngitis is typically treated with antibiotics; it can lead to suppurative complications (acute otitis media, sinusitis, retro- and peritonsillar abscess, cervical adenitis), rheumatic fever, and organ damage if left undiagnosed and untreated.^3^

The gold standard method for diagnosis of GAS pharyngitis is swabbing the posterior pharynx and testing with a rapid antigen detection test (RADT); standard culture is performed if the RADT is negative.^4, 5^ However, culture is time-consuming and may delay the diagnosis.^6^ Currently, there are numerous advanced techniques for the detection of *S. pyogenes*, such as polymerase chain reaction (PCR),^7, 8^ multiplex PCR,^9^ impedimetric biosensors,^10, 11^ touch spray-mass spectrometry,^12^ and commercial test kits (e.g., BinaxNOW® Strep A Test; OSOM® Ultra Strep A Test; BD Chek™ Group A Strep; QuickVue In-line Strep A Test). Some commercially available at-home GAS test kits, such as ezlevel Strep A test, © ezleveltests, rely on the parent swabbing the child’s throat; this is not the standard of care and it does not produce reliable results.^13^ Even in a clinical setting it can be challenging to obtain a high-quality swab sample of the posterior pharynx including the tonsils due to child resistance and discomfort.^14^ Children may even refuse to use or be noncompliant with oral sample collection, which can lead to skewed results.^14^ This problem is exacerbated in very young children and those with developmental disabilities. Recently, saliva samples have been shown to be an alternative to pharyngeal swabs for respiratory disease diagnosis.^15, 16^ A recent study of GAS in the saliva of children used swabs to absorb saliva and PCR for detection; 19 of 20 patients with previously confirmed GAS pharyngitis tested positive.^16^ In the present work, we aim to further improve the patient experience and establish a salivary sampling open-fluidic device for *S. pyogenes* capture, which can be used comfortably in a clinic or home setting for adults and children.

Our open-fluidic sampling device, CandyCollect, is inspired by a lollipop; it includes a small polystyrene stick with one end coated in isomalt and fruit flavoring candy. We aimed to engineer a device that is both efficient at capturing bacteria in saliva samples and decreases the discomfort experienced in traditional swabbing methods. We demonstrate that the mass of the candy can be used to inform the sampling time, effectively using the disappearance of the flavoring as a built-in timer. Additionally, we conducted *in vitro* experiments involving the target pathogen, *S. pyogenes*, to investigate its adherence to the device, the effect of sample evaporation on elution and the ability to detect bacteria on the devices using quantitative polymerase chain reaction (qPCR). We have successfully demonstrated that the open-fluidic channel prevents the tongue from scraping the surface and retains the bacteria within the channel, demonstrating that our device can capture the pathogen and accumulate more bacteria over time without disturbance from external factors. To assess the usability and feasibility of this device, we sent CandyCollects to 17 participants across the country to record the person-to-person differences in candy dissolving time and user feedback. With the growing need for decentralized medicine and telehealth solutions, our device provides an accessible sampling method for more versatility in at-home and clinical settings.

## Methods

### CandyCollect fabrication

CandyCollect sticks were fabricated out of 2 mm thick polystyrene sheets (Goodfellow, Cat# 235-756-86), and an acrylic mold for making the silicone mold was fabricated from a 25.4 mm thick acrylic (Source One) by CNC milling (Datron Neo). Dimensions of the CandyCollect stick are provided in Figure S1. The silicone mold used to cast the CandyCollect was made by pouring food safe silicone (smooth-sil 940, Smooth-On) into the acrylic mold, and cured for 24 hrs at room temperature. The candy was prepared following hygiene guidance from the Washington State Cottage Food Operations Law (RCW 69.22.040(2b-f(ii-iv)). The study team member who prepared the CandyCollect trained in food safety, has a Food Worker Card, and wore a mask during food preparation. In brief, the candy was prepared by gradually adding 1000 g of isomalt to 80 g of boiling water until the solution reached ∼170°C. Then 3.5 mL of Chefmaster Super red gel color was added after the solution reached 165°C. Then 10 mL of LorAnn Super Strength Strawberry Candy Oil (LorAnn, Cat# 0320-0500) was added after the solution reached 170°C, and it was removed from the heat. The candy was cooled to room temperature and remelted in small portions for distribution among molds. The silicone mold was placed onto a marble slab to ensure rapid cooling, then the melted candy was poured into each individual CandyCollect shape until the CandyCollect was filled. A CandyCollect stick was placed into the candy, channels facing up, with the circular portion of the polystyrene centered in the candy. The stick was held in place for 15 s, then the next stick was placed. The sticks were pushed back into the candy as needed (using sterilized metal tools) as they set for approximately 5-10 min. Once set, they were carefully removed from the silicone mold and placed on a silicone mat on the marble to finish cooling. After cooling, the CandyCollect were placed into oriented polypropylene bags (3×4 inch BakeBaking) and heat sealed using an impulse sealer (Metronic-Model: FS-200) on setting 3.

### CandyCollect dissolving time in human subjects

*Participant characteristics*. This study was approved by the University of Washington Institutional Review Board (IRB) under IRB-exempt protocol STUDY00012318. All study procedures were performed after informed consent was obtained. A total of 17 healthy volunteers over the age of 18 years were recruited via the University of Washington Institute of Translational Health Sciences (ITHS) “participate in research” website and the study team website.

#### Inclusion and exclusion criteria for participant enrollment

Inclusion criterion: over the age of 18 years. Exclusion criteria: individuals who are allergic to sugar alcohols or are in vulnerable groups including pregnant women or individuals residing in a correctional facility.

#### Enrollment of participants

Once potential participants responded with an email sharing their interest in participating, a study team member sent them a link to a pre-screening survey which asked questions about the inclusion and exclusion criteria. Eligible participants were then asked to enter their demographic information, their first and last name, shipping address, and contact information. After signing an informed consent form on REDCap (Research Electronic Data Capture, hosted at the ITHS),^17^ participants were enrolled into the study. All identifiable data were securely stored in REDCap and only de-identifiable data were obtained for further analysis.

#### Human subjects study procedures

Study participants were enrolled in order to test the general feasibility of the CandyCollects as a collection method. A kit containing six CandyCollect devices and an instruction card was sent to each participant. As directed on the instruction card, each participant consumed one CandyCollect at a time and recorded the total time it took to dissolve the candy on each device. Study data were collected and managed using REDCap.

### *S. pyogenes* culture and imaging

#### Todd-Hewitt broth (TH broth) liquid media preparation

For the THY liquid media, 30 g of Todd-Hewitt Broth (BD Bacto™ TH broth, Fisher Scientific, Cat# DF0492-17-6) and 2 g of Yeast Extract (United States Biological Corporation, Fisher Scientific, Cat# NC9796728) (THY) were added to 0.8 L distilled water and dissolved to completion. Additional distilled water was added for a total volume of 1 L. THY liquid media was autoclaved for 30 min, cooled to room temperature and stored at 4°C.

#### S. pyogenes maintenance in agar plate

The *S. pyogenes* was prepared from *Streptococcus pyogenes* Rosenbach (American Type Culture Collection, ATCC^®^, Cat# 700294™). Freeze-dried *S. pyogenes* was wetted with 1 mL liquid media, and then transferred to another conical tube containing 4.4 mL of THY liquid media. To maintain and ensure isolated colonies, *S. pyogenes* was cultured on Trypticase™ Soy Agar (TSA II™) with Sheep Blood (BD BBL™, Fisher Scientific, Cat# B21239X). A sterile disposable inoculating loop (Globe Scientific, Fisher Scientific, Cat# 22-170-201) was dipped in liquid media containing *S. pyogenes* and streaked along the edge of the agar plate. The plate was then rotated 90° and the streaking was continued. This was repeated two times more using the same loop. The agar plate was maintained in an incubator at 37 °C with 5% carbon dioxide overnight, then stored at room temperature for up to seven days.

#### Incubation of S. pyogenes in liquid media

To ensure a pure culture, fresh *S. pyogenes* from agar plates were inoculated in liquid media one day before the experiment. A pipette tip was used to pick one colony from the agar plate containing viable *S. pyogenes* colonies, and then the pipette tip was stirred in 7 mL THY liquid media in a sterile 14 mL snap-cap round bottom polystyrene tube (Falcon®, Corning, Product# 352001). *S. pyogenes* was incubated at 37 °C with 5% carbon dioxide in the incubator overnight.

#### Capturing, fixing and staining of S. pyogenes

The concentration of *S. pyogenes* was measured through Visible 721-Vis Spectrophotometer (vinmax) at a wavelength of 600 nm. *S. pyogenes* in liquid media was centrifuged in 1 mL at 10,000 rpm for 10 minutes. 50 µL of *S. pyogenes* suspension at the concentration of 1×10^9^ CFU/mL was applied to each CandyCollect device and then incubated for 10 minutes (unless otherwise specified). The device was rinsed with PBS and fixed with 4% paraformaldehyde (PFA) for 15 minutes. To remove excess PFA, the device was rinsed with PBS. 50 µL of Alexa Fluor™ 488 Wheat Germ Agglutinin (WGA, Invitrogen™, Fisher Scientific, Cat# W11261, 1 mg/mL) at 1: 500 dilution (v/v) was added to the channel and incubated for 30 minutes at room temperature in the absence of light. Alexa Fluor™ 488 conjugated WGA was used to stain bacteria cell wall peptidoglycans. Bacteria was green fluorescently labeled after staining. The device was rinsed with PBS to remove excess WGA. Then the device was stored in centrifuge tubes containing PBS.

#### Imaging and quantification

Fluorescent images of *S. pyogenes* were obtained on a Zeiss Axiovert 200 with a 10× (0.30 NA) magnification coupled with Axiocam 503 mono camera (Carl Zeiss AG, Oberkochen, Germany). Multiple channel regions (four to five) were chosen randomly from each device to take images for confirming no bias on a specific region. For all images, the contrast was adjusted uniformly using Fiji (ImageJ) software. To measure the integrated density, each image was further processed using functions from ImageJ. We used the modified procedure from Theberge *et al*., 2015.^18^ Specifically, each image was converted to 8 bits. Then the background was subtracted using “Subtract Background”. Next, to convert the images into black and white, a default threshold using the “Li Dark” function was used. Three regions of interest (ROI, 200 × 200 µm^2^) were selected to measure the integrated density (Fig S2).

#### Statistics

Statistical analysis was performed using GraphPad Prism 9 software. One-way Analysis of Variance (One-way ANOVA) was chosen to compare groups and Tukey’s multiple comparison tests were further used in evaluating significance of pairwise comparisons.

### Cotton swab scraping assay

50 µl of *S. pyogenes* suspension at the concentration of 1×10^9^ CFU/mL were applied on each CandyCollect device (Figure 2B top panel) and the device without channels which contained a milled shallow circle (Figure 2B bottom panel) and incubated for 10 min. The devices were rinsed once with PBS and excess liquid was drained with KimWipes. Strokes were made on the surface of each device using a cotton swab, to mimic scraping by the tongue. Control devices were not touched by cotton swabs. To evaluate the effect of cotton swab scraping on bacteria binding, the bacteria on the device were stained and imaged following the protocol stated above.

### Plasma treatment shelf life experiments

*Oxygen plasma treatment*. The devices were plasma treated by Zepto LC PC Plasma Treater (Diener Electronic GmbH, Ebhausen, Germany). The chamber was pumped down to a pressure of 0.20 mbar, gas was supplied (0.25 mbar for 2 minutes), power enabled (70W for 5 minutes). After plasma treatment, the devices were stored in OmniTrays (Thermo Scientific™ Nunc™ OmniTray™, Fisher Scientific, Cat# 12-565-296) under room temperature for zero day (control group), three days, one week, two weeks, two months with n=3 replicated devices before the experiment. For time points less than one month, devices were treated in descending order (i.e., two-week devices treated first, followed by one-week devices 7 days later) allowing for the two-week, one-week, and three-day devices to be tested on the same day. The two-month devices were tested on a different day. The zero day (control group) was done for every experiment set. The bacteria on the device were incubated, stained, and imaged following protocol stated above.

### Saliva and candy interference experiment

A stock solution of isomalt was made at a 1.0 g/mL concentration in phosphate buffered saline (PBS). This solution was mixed with pooled saliva (Innovative Research Inc., Fisher Scientific, Cat# 50-203-6057) in a one-to-one ratio (final isomalt concentration 0.5 g/mL). This process was repeated with the device candy (composition: isomalt, water, candy oil flavoring, and gel food coloring). These solutions were used to resuspend *S. pyogenes* at a concentration of 1.7×10^9^ CFU/mL.

### Isolation and purification of genomic DNA from *S. pyogenes*

DNA was isolated using DNeasy UltraClean Microbial Kit (QIAGEN, Cat# 12224) according to the protocol supplied by the manufacturer. *S. pyogenes* was cultured overnight in THY broth. Bacterial cells were pelleted and resuspended in PowerBead Solution. 300 µL of bacteria suspension containing approximately 2×10^9^ CFU of *S. pyogenes* was transferred to each PowerBead Tube followed by the addition of 50 µL of Solution SL. The tubes were placed in a MiniBeadBeater (BioSpec Products, Bartlesville, OK USA) and bead-beating was applied twice for 30 seconds as an alternative to vortexing to facilitate cell lysis. The lysates were loaded onto MB spin column. Then DNA was eluted from each column with 50 µL of 10 mM Tris-HCl buffer. DNA concentrations were measured using BioTek Cytation 5 Cell Imaging Multi-mode Reader (Agilent, 300 Industry Drive, Pittsburgh, PA).

### Elution of *S. pyogenes* from CandyCollect devices and preparation of cell lysates containing bacterial genomic DNA

The devices were incubated with 50 µL of *S. pyogenes* suspension for 10 min at the following concentrations: 1×10^3^, 1×10^4^, 1×10^5^, 1×10^7^, 1×10^9^ CFU/mL. Devices eluted immediately, before the suspension dried, were defined as “wet samples”. Devices placed in 15 mL tubes and allowed to dry for 24 hours inside a biosafety cabinet were defined as “dry samples”; for these samples the following concentrations were used: 500, 1×10^3^, 1×10^4^, 1×10^5^CFU/mL of *S. pyogenes*. All devices were eluted in 300 µL of ESwab^™^ buffer (Becton, Dickinson and Company, Cat# R723482) with 5% ethanol in 15 mL round bottom tubes that accommodate the CandyCollect device. To facilitate complete removal of bacteria from devices, ∼ 50 µL of 0.1 mm Zirconia/Silica beads (BioSpec Products, Cat# 11079101Z) was added to each tube and the tubes were vortexed for 50 seconds to dislodge the captured bacteria on the devices. In addition, the tubes were heated at 85 °C for 10 min. The resulting bacteria suspension containing beads was transferred to a screw cap microtube (ThermoFisher, Cat# 3490). Bead beating was carried out on the above-mentioned BeadBeater twice for 30 sec to aid the release of DNA from bacteria. Genomic DNA in bacterial cell lysates was used as template in the qPCR assay (This protocol is used for Figure S3).

### DNA enrichment

Genomic DNA in bacterial cell lysates was enriched using Invitrogen™ ChargeSwitch™ gDNA Mini Bacteria Kit (Fisher Scientific, Cat# CS11301). Following the protocol supplied by the manufacturer, genomic DNA was concentrated 10 times. In brief, genomic DNA adheres on the surface of the magnetic beads when the buffer pH is lower than 6.5. Then the adhered DNA is eluted off of the beads by the provided elution buffer. Concentrated genomic DNA in bacterial cell lysates was also used as template in the qPCR assay (This enrichment procedure is used for Figure 4C).

### Quantitative PCR assay for detection of *S. pyogenes*

A putative *S. pyogenes* species-specific transcription regulator, *spy1258* (GenBank accession No. AE006565), was used for detecting *S. pyogenes*. The primers/probe targeting this gene have been used in identification and/or quantification of *S. pyogenes* previously.^19, 20^ The primers/probe sequences designed by CDC Streptococcus Laboratory^20^ were adapted for qPCR detection of *S. pyogenes* in our assay: the forward primer: 5’-GCA CTC GCT ACT ATT TCT TAC CTC AA-3’; the reverse primer: 5’-GTC ACA ATG TCT TGG AAA CCA GTA AT-3’; the probe sequence: 5’-FAM-CCG CAA C”T”C ATC AAG GAT TTC TGT TAC CA-3’-SpC6, “T” = BHQ1. The primers were obtained from IDT (Integrated DNA Technologies, Inc., Coralville, IA, USA). The probe was ordered from MilliporeSigma (Summit Drive Burlington, MA USA). PerfeCTa® qPCR ToughMix (VWR, Cat# 97065-954) was used with 10 µL of DNA template in the qPCR assay with a total volume of 25 µL per reaction. The final concentrations of both forward and reverse primers were 300 nM; the probe concentration was 100 nM (Figure 4B, 4C, and S4B). For select experiments, qPCR analysis was processed with primer concentration 500 nM, and probe concentration 250 nM (Figure S3Ai and S3Aii). CFX connect Real-Time PCR Detection System (Bio-rad Laboratories, Hercules, CA, USA) was used to perform PCR and collect fluorescence data during DNA amplification. The qPCR runs 40 cycles at 95 °C for 15 s and 60 °C for 30 s after initial denaturation of 95 °C for 5 min. Data analysis was carried out using CFX Maestro software version 2.2. 1:10 serial dilutions of purified genomic DNA ranging from 50 ng to 50 fg were used as standards. PCR was performed in 96-well PCR plates (Bio-Rad Laboratories, Hercules, CA, USA) in either duplicate or triplicate.

## Results and Discussion

### Design of the CandyCollect device

Noncompliance with diagnostic testing in children affects accuracy and validity of diagnosis for upper respiratory infections, and the inability to access a medical professional also prevents children from receiving appropriate diagnosis and treatment. When left untreated, infections such as strep throat can have severe consequences such as rheumatic fever, organ damage, secondary infections, and death.^1, 2, 21^ To optimize the diagnosis of streptococcal pharyngeal infections we have developed a saliva sampling device that takes design inspiration from a classic childhood treat, a lollipop. CandyCollect was designed to comfortably fit in a child’s mouth, mitigate any potential choking hazard by use of a thin candy coating rather than a large piece of candy, be compatible for use with common elution kits, such as ESwab 480C (Copan), and have optimal saliva exchange with the capture surface. The CandyCollect sampling device consists of a computer numerical control (CNC) milled polystyrene stick that is coated on the back and edges with isomalt candy. The face of the stick is not coated in candy and contains a plasma treated open-fluidic spiral channel for saliva and bacteria capture (Figure 1). Plasma treatment is known to facilitate bacterial adhesion on polystyrene surfaces.^22, 23^ Oxygen plasma treatment increases the substrate’s hydrophilicity and aqueous wettability resulting in greater bacterial adhesion.^23^ The aspect ratio of the channel was designed such that a tongue cannot scrape the bottom of the channel and remove captured bacteria. Isomalt, a sugar alcohol, was used for the candy because it has a low hygroscopicity (ERH of 88), meaning it will not absorb water until equilibrium relative humidity reaches 88% at 20°C,^24^ which enables it to remain stable if packaged and stored for extended periods of time. Additionally, in the presence of oral bacteria, such as *Streptococcus mutans*, sucrose ferments and becomes turbid while isomalt and other polyols do not.^24^ Finally, isomalt has a glycemic index of two and is safe for diabetic patients.^25^ The candy also acts as a built-in timer for sampling time; it ensures the patient keeps the device in their mouth for the optimal duration to achieve capture of bacteria. When the candy flavoring disappears, the sampling is complete, and the patient can mail the CandyCollect back or schedule a pickup for analysis. By varying the amount of isomalt on the CandyCollect we can adjust for different sampling times required by other respiratory pathogens. The ability to enhance or accumulate the pathogens of interest over time shows the future potential of CandyCollect to capture a range of pathogens that are present at different concentrations. Determining how much *S. pyogenes* the device can capture is important in determining if the device will be compatible with current quantification or rapid PCR methods such as qPCR or real-time PCR.

**Figure 1.**
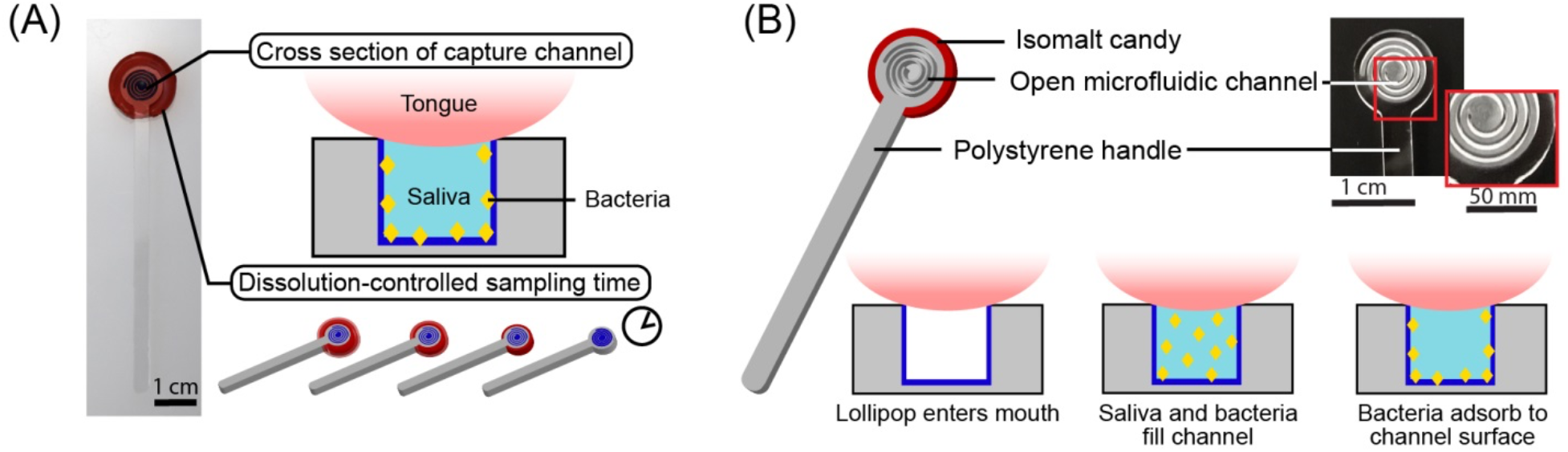
The CandyCollect is a lollipop-inspired, at-home, saliva collection and bacteria sampling device for *S. pyogenes*, the bacteria that causes pharyngeal GAS infection (commonly called strep throat). (A) The open-fluidic channel in the CandyCollect captures the bacteria from saliva, and the candy flavoring is a built-in timer for sampling time (i.e., dissolving time of the candy). The open-fluidic channel is designed to prevent the tongue from scraping the bottom of the channel and removing the collected bacteria, allowing bacteria to accumulate during the sampling time. (B) The CandyCollect is composed of a polystyrene stick with a microfluidic channel and red isomalt candy. When the CandyCollect enters the mouth the channel fills with saliva and the bacteria adheres to the plasma treated surface of the channel.

### CandyCollect device enables bacteria accumulation over time and prevents loss due to mechanical scraping

Additionally, determining if bacteria collection increases over time relies on tailoring the mass and surface area ratio of the candy to control the sampling time. *In vitro*, our device can accumulate *S. pyogenes* over time and capture it in as little as 30 seconds; longer incubation time increases the amount of bacteria accumulated (Figure 2Ai and 2Aii). Further, it was important to show that the channels prevented the tongue from scraping off any captured *S. pyogenes* (Figure 2B). A shallow circle was milled for the flat surface of the CandyCollect device. Fluorescence microscopy images indicate the channel protects the *S. pyogenes* from being scraped off as opposed to the flat surface which shows a marked decrease in *S. pyogenes* after scraping (Figure 2B).

**Figure 2.**
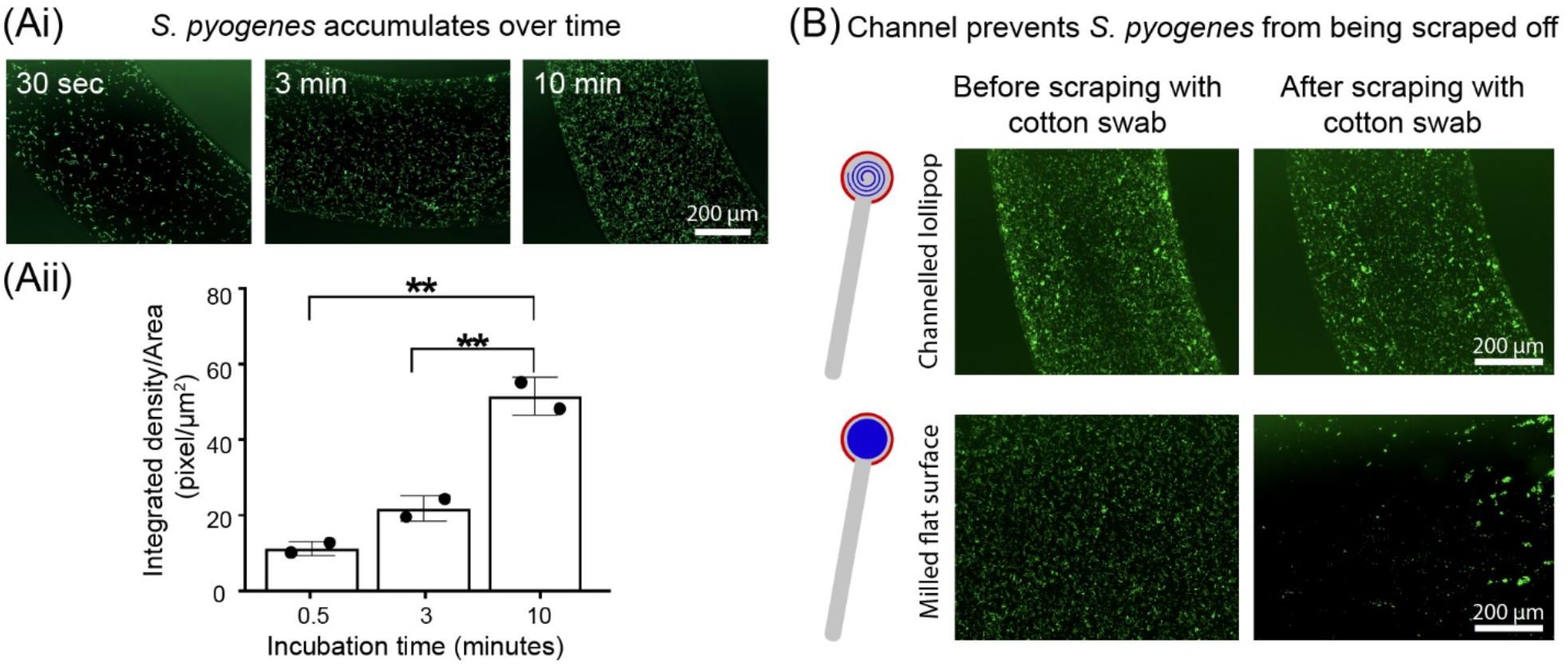
*S. pyogenes* accumulates over time, and the channel geometry prevents loss of captured *S. pyogenes*. (Ai) *S. pyogenes* captured *in vitro* on the CandyCollect accumulates over 30 sec, 3 min, and 10 min. (Aii) Quantification of the integrated density per area (pixel/µm^2^). Each data point represents an individual CandyCollect; The bar graph represents the mean ± SD of n = 2 CandyCollects. Data sets were analyzed using one-way ANOVA; p-values are indicated for pairwise comparisons between different incubation time: \*\**p* ≤ 0.01 (Tukey’s multiple comparison tests). (B) *S. pyogenes in vitro* remains in the channel following scraping with a cotton swab (intended to represent the tongue scraping the device) compared to markedly reduced amounts in a CandyCollect without a channel (milled flat surface) at the same concentration. Note: an incubation time of 10 min was used in B. Images are representative of 3 independent experiments, with duplicate devices and 4 images taken per replicate, and the data point plotted is the average. *S. pyogenes* was green fluorescently labeled.

### Shelf life tests: CandyCollect effectively captures *S. pyogenes* after 2 months of storage

Our device was designed to enable a high-quality, comfortable collection of pathogens from saliva in a home or clinical setting. We anticipate the devices to be stored in homes or clinics, thus it is highly likely the devices would be stored for extended periods of time before usage. It is known that over time plasma treated surfaces become more hydrophobic (water contact angle increases within 10 days and stabilizes with long storage time).^26^ To test whether the time-induced changes in the plasma treated surface of CandyCollect affect its ability to capture bacteria, we conducted shelf life experiments at room temperature. Fluorescence microscopy images show that there is no significant difference observed in the images between 0 day (control group), 3 days, 1 week, and 2 weeks (Figure 3Ai and Aii). An additional experiment was conducted to test two months of storage, which also showed the device was still able to capture *S. pyogenes* efficiently compared to the control (Figure 3Bi and Bii). In the future, we plan to conduct shelf life experiments for lengths such as six months to one year.

**Figure 3.**
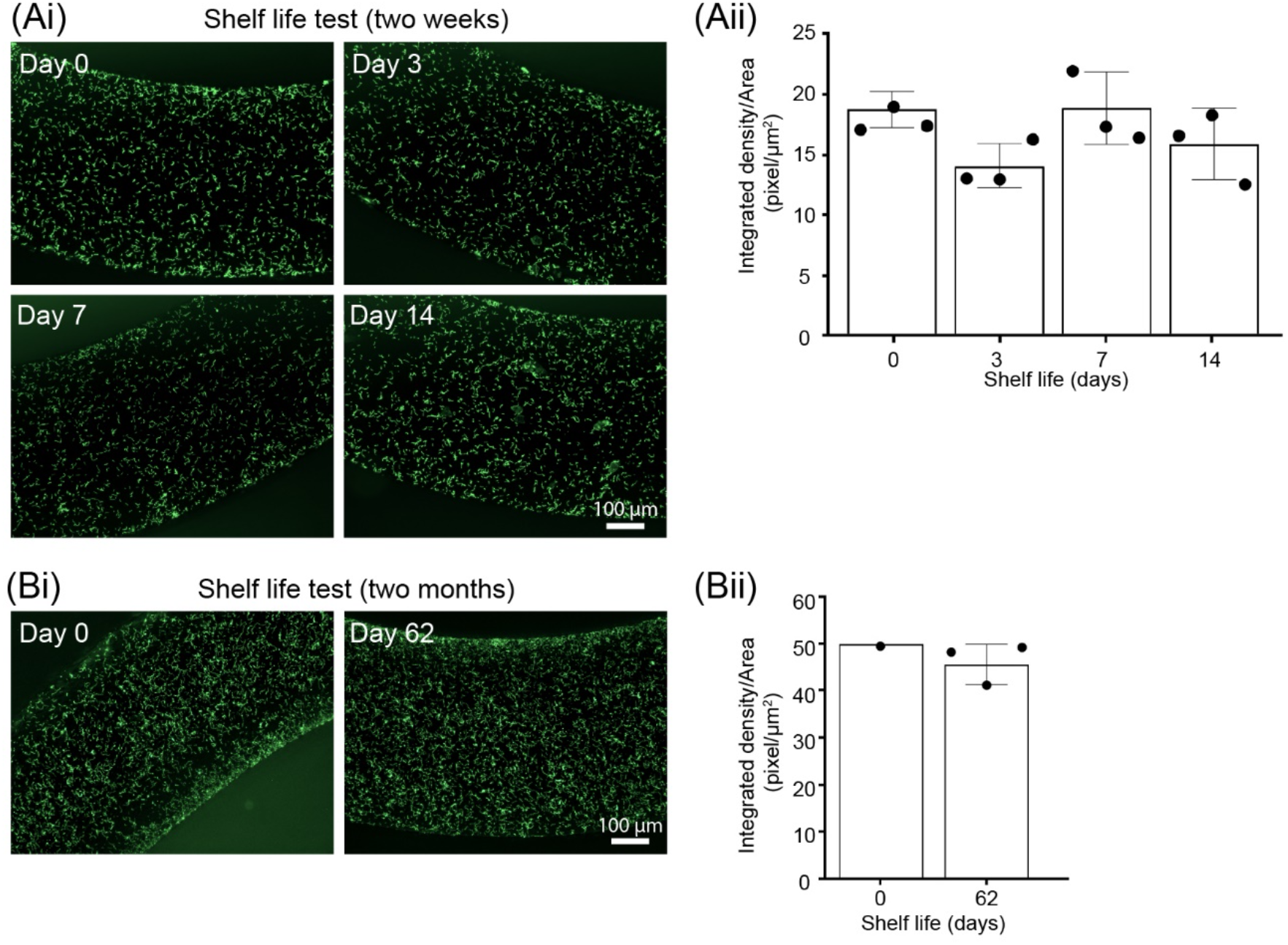
Shelf life tests demonstrate that CandyCollect effectively captures *S. pyogenes* after 2 months of storage. Devices were plasma treated and stored at room temperature for (Ai) 0 days (control group), 3 days, 1 week, 2 weeks. Fluorescence microscopy images indicate capture of *S. pyogenes* after 14 days of storage is similar to the control. (Aii) Quantification of the integrated density per area (pixel/µm^2^). Data sets were analyzed using one-way ANOVA; no significant difference between shelf life periods was found. (Bi) 0 day (control group) and 2 months. Fluorescence microscopy images indicate capture of *S. pyogenes* after 62 days (∼2 months) of storage is similar to the control. (Bii) Quantification of the integrated density per area (pixel/µm^2^). Note: in both A and B, *S. pyogenes* at a concentration of 1×10^9^ CFU/mL was incubated on the CandyCollect device for 10 minutes. Each data point represents an individual CandyCollect device (4 images were taken per device, and the data point plotted is the average); the bars represent the mean ± SD of n=3 CandyCollects. *S. pyogenes* was green fluorescently labeled.

### Elution of *S. pyogenes* from CandyCollect devices and qPCR detection

The saliva on the devices will evaporate in the time between sampling and analysis if they are shipped back to the laboratory. It is important to note that we will not be culturing the bacteria from the CandyCollect device but rather eluting the bacteria from the device for qPCR analysis to detect the presence or absence of *S. pyogenes*. To evaluate whether drying affects the ability to elute *S. pyogenes*, we left the devices at room temperature overnight to evaporate. Fluorescence microscopy images showed that the *S. pyogenes* captured on devices could still be eluted from the device for further experiments (Figure 4A).

**Figure 4.**
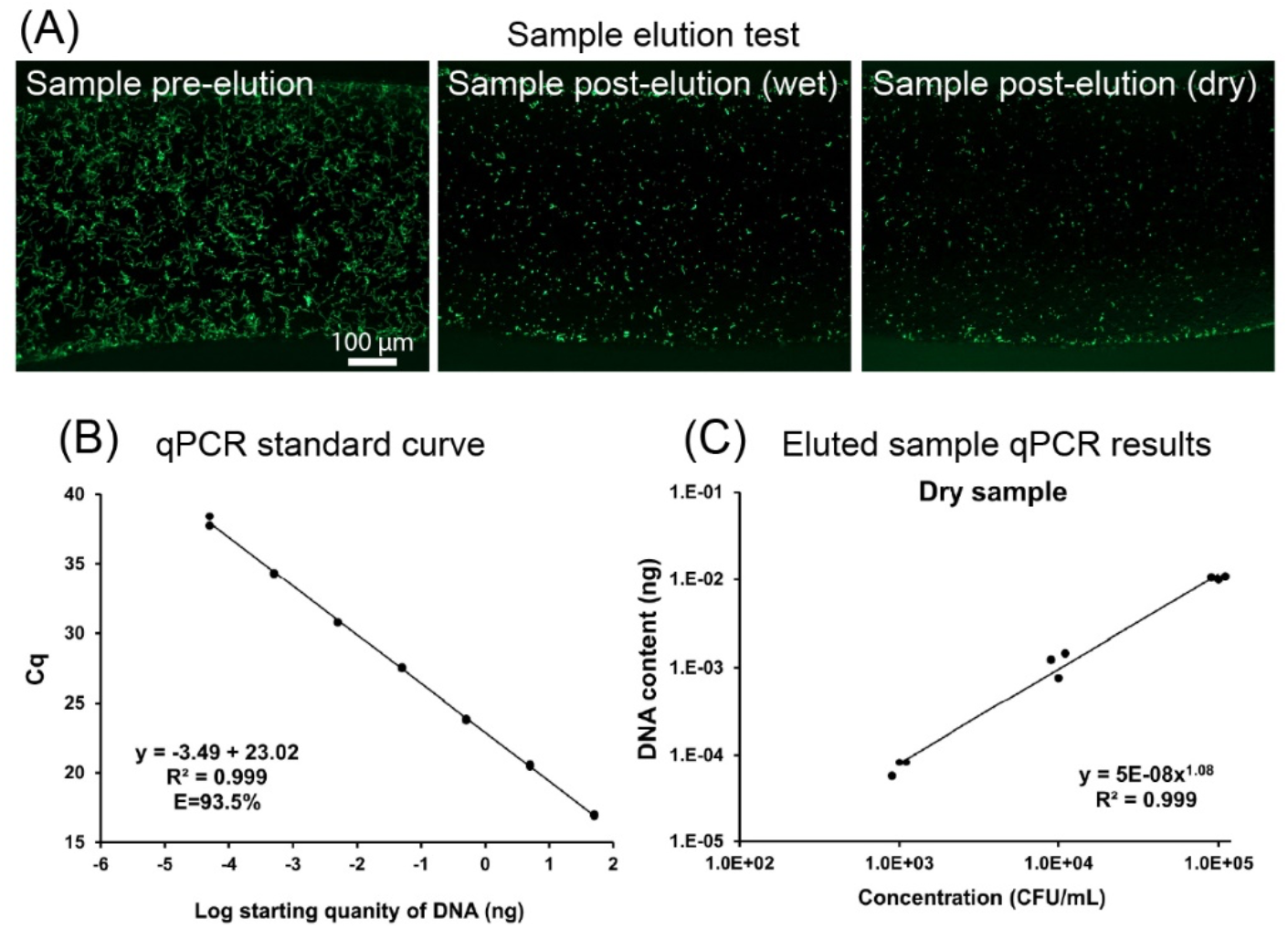
Elution tests demonstrate that *S. pyogenes* can be eluted from the CandyCollect and analyzed by qPCR. (4A) Elution of *S. pyogenes* is successful in wet and dry samples. *S. pyogenes* at a concentration of 1×10^9^ CFU/mL was incubated on the CandyCollect device for 10 minutes. *S. pyogenes* was green fluorescently labeled. The image results suggest that the *S. pyogenes* can be eluted from CandyCollect device and drying does not hinder bacteria elution. (4B) Standard curve for the qPCR assay. 1:10 serial dilutions of genomic DNA ranging from 50 ng to 50 fg were used as template for qPCR. Each dot represents one technical duplicate (in cases where one point is visible the duplicates were identical). The standard curve in which Cq values were plotted against starting template DNA, was linear from 50 ng to 50 fg. qPCR efficiency ranged from 93.5-100.6% across four independent experiments. (4C) Device drying did not prevent elution and detection of *S. pyogenes* sampled on devices. Quantification of *S. pyogenes* by qPCR. Each data point represents an individual CandyCollect. Three concentrations of *S. pyogenes* suspension (1×10^3^, 1×10^4^, 1×10^5^ CFU/ml) were used for the experiment. DNA content was detected in a bacterial concentration-dependent manner. Devices run through the incubation and elution protocols without bacteria loading were used as device negative controls. No-template controls (NTC) for qPCR and device negative controls had no detected signal.

We also developed an elution method to efficiently remove bacteria from the devices for downstream analysis. Furthermore, we successfully established a qPCR assay to detect *S. pyogenes* eluted from CandyCollect devices (Figure 4B and C). The assay can determine the number of bacteria by detection of genomic DNA content. DNA content measured in the bacterial samples eluted from the devices by qPCR was highly correlated with bacterial concentrations that were incubated on the device (R^2^=0.991-1, Figure 4C and S3). Importantly, a single DNA product was amplified when using DNA templates from *S. pyogenes*, but no products were detected when using templates from either *Streptococcus mutans* or *Staphylococcus aureus* (Figure S4), demonstrating high selectivity in detecting *S. pyogenes*. In addition, our assay standard curve showed linearity down to 50 fg (Figure 4B), representing approximately 25 CFU of *S. pyogenes*, demonstrating the ability to detect low levels of *S. pyogenes* (Figure 4B). Our results showed DNA was detectable at bacterial concentrations of 1×10^5^, 1×10^4^, and 1×10^3^ CFU/mL in both “dry samples” and “wet samples” (Figure 4C and S3). (Refer to the methods section for definitions of dry and wet samples.) *S. pyogenes* concentrations *in vivo* range from 500 to 1.5×10^4^ CFU/mL from pharyngeal samples in patients with streptococcal pharyngitis.^27^ These results suggest that drying does not hinder bacterial elution from CandyCollect device nor affect the ability to detect DNA at a bacterial concentration of 10^3^ CFU/mL. We will further develop the extraction protocol and qPCR assay in future work to optimize the limits of detection of our device.

The pathogens found in human saliva will have different typical concentrations, therefore, flexibility in dissolving time of the candy is desirable. To demonstrate flexibility in dissolving time, six CandyCollects with different masses of candy were created to determine how the mass and dimensions of the candy affect the dissolving time. One individual consumed three CandyCollects of each version and recorded the dissolving time (Figure 5A). Based on the results, the CandyCollects dissolved between less than one minute and nearly fifteen minutes, accommodating a range of sampling times that may be needed depending on the target bacteria. The full set of data including the mass and dimensions of the CandyCollects is available in Table S1. However, the CandyCollects are not limited to this range and can be made larger to accommodate sampling times longer than fifteen minutes.

**Figure 5.**
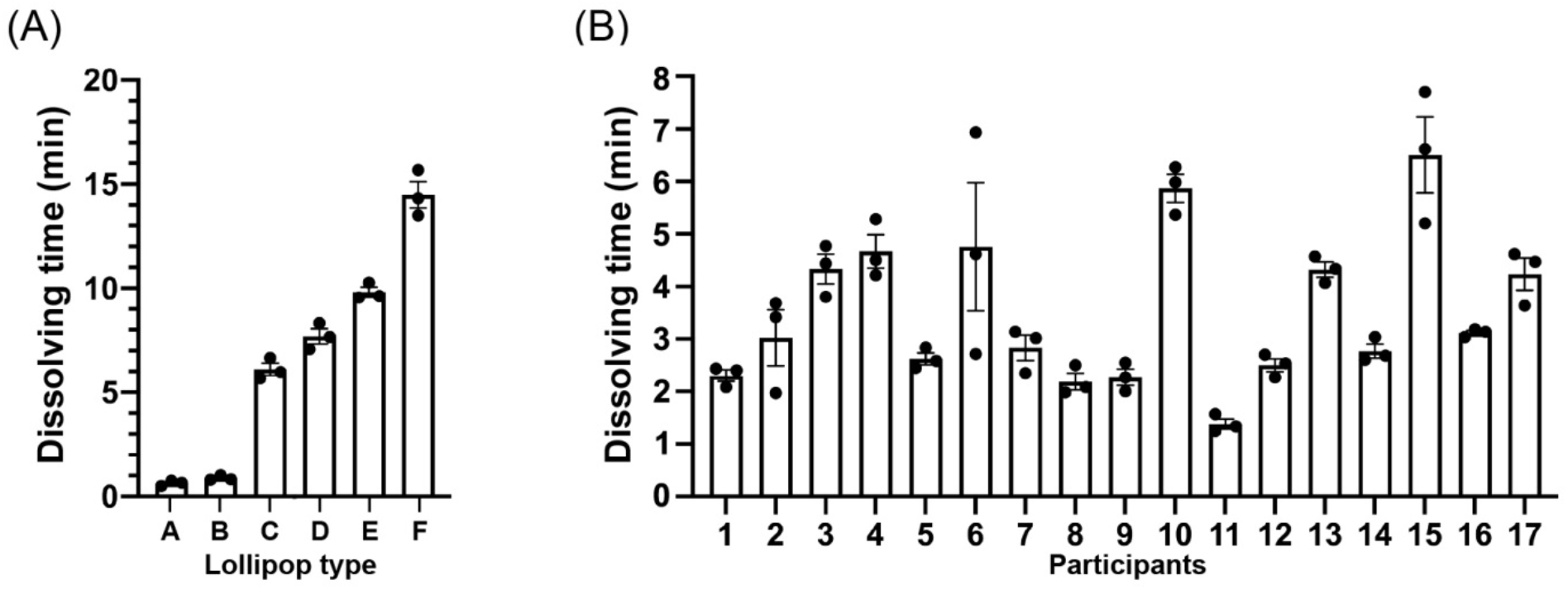
CandyCollect dissolving time can be controlled by changing the mass and size of the CandyCollect. (A) Six types of CandyCollect devices, each with a different mass and size of candy, were consumed by one individual. Bar graphs represent mean ± SEM of n = 3 CandyCollects. Please refer to Table S1 in the Supporting Information for CandyCollect mass and dimensions. (B) A single size of CandyCollect devices was sent to 17 research participants for a usability test. The mass of the CandyCollect in B is 0.90-1.10 g, with a diameter of 16 mm, and thickness of 4 mm. Bar graphs represent mean ± SEM of n = 3 CandyCollects.

To investigate the variability in dissolving time across different people, we enrolled participants in a pilot feasibility and usability study. The participants were instructed to consume three CandyCollects that were 0.90-1.10 g, with a diameter of 16 mm, and a thickness of 4 mm and record the dissolving time (Figure 5B). The average dissolving time across participants is 3.51 mins, with a minimum of 1.25 mins; the majority of participants completely consumed the candy within 2-4 mins. The full set of data is available in Table S2. In practice, the required sampling time, which depends on the abundance of a pathogen of interest and the threshold of detection by qPCR, will be evaluated for each pathogen of interest, and the mass of the candy will be adjusted accordingly. We understand that there is personal variability in the amount of time required for the candy to dissolve, and we will adjust the mass and dimensions of the candy so that the minimum dissolving time matches the required collection time for a given pathogen.

During the collection period, the device channel will encounter saliva, bacteria (the analyte of interest) and candy. Isomalt, the primary component of the candy fixed to the device, has been demonstrated to have little impact on bacterial proliferation,^24^ however the candy, in addition to saliva, both have the potential to interfere with the adhesion of *S. pyogenes* to the channels of the device, either through competitive binding or chemical action. To replicate the environment of the device during sample capture in a participant, *S. pyogenes* was incubated *in vitro* on the device in mixtures containing pooled saliva, isomalt and/or candy (Figure 6). Stock solutions of isomalt and the device candy were made in phosphate buffered saline (PBS). These solutions were mixed with pooled saliva in a one-to-one ratio for a final concentration of isomalt and candy of 0.5 g/mL in their respective solutions in order to maximize the concentration of the isomalt and candy in the saliva. Solutions were used to resuspend *S. pyogenes* and subsequently applied to the devices and incubated. Fluorescence images demonstrate that saliva, isomalt, and the device candy do not impact the adhesion of *S. pyogenes* to the channel of the device (Figure 6).

**Figure 6.**
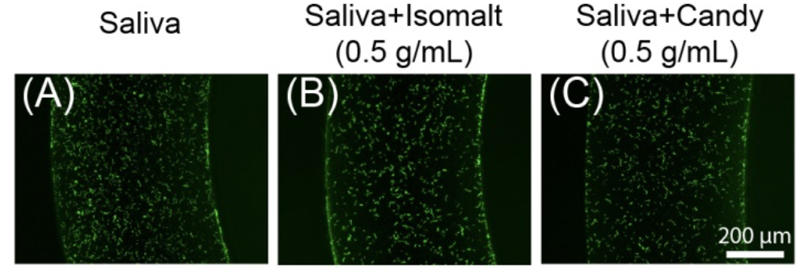
Saliva, isomalt, and other candy ingredients do not interfere with *S. pyogenes* capture on the CandyCollect device. *S. pyogenes* was green fluorescently labeled. Fluorescence microscopy images of captured *S. pyogenes* in the device when incubated in solutions containing (A) pooled saliva, (B) pooled saliva and isomalt, and (C) pooled saliva and device candy (composition: isomalt, water, candy oil flavoring, and gel food coloring). Solutions were used to resuspend *S. pyogenes* at a concentration of 1.7×10^9^ CFU/mL, incubated on the CandyCollect device for 2 minutes, and imaged. Images are representative of one independent experiment, with 2 replicates and 3-4 images taken per replicate.

## Conclusion

In this work, we developed a saliva sampling device that aims to enable high-quality, comfortable collection of pathogens from saliva in a home or clinical setting. **This work shows (1) the CandyCollect device can effectively capture and concentrate bacteria for further analysis, (2) the CandyCollect device is functional after storage times of up to two months, (3) candy flavoring can be used as an indicator for sampling time to facilitate effective bacterial capture, and (4) qPCR can be successfully performed on samples eluted from the CandyCollect device**. In the future, manufacturing of the proposed device can be scaled up using rapid injection molding and testing of the CandyCollect in a clinical setting will provide useful information as to the widespread utility of this device. This technology has the potential to bring sampling technologies into decentralized telemedicine and minimize the need to access a clinic. Furthermore, this CandyCollect platform and advanced sample analysis has great potential for sample collection and diagnosis in other respiratory diseases. Testing of the CandyCollect in a clinical setting will provide useful information as to the widespread utility of this device.

## Acknowledgements

This work was supported by a Washington Research Foundation (WRF) Phase 1 Grant, National Institutes of Health grants (R21AI147502-02, R35GM128648 (for some of the in-lab developments and in vitro experiments)), the Society for Laboratory Automation and Screening (SLASFG2020, UNL), the David and Lucile Packard Foundation, an Alfred P. Sloan Research Fellowship, and the University of Washington. REDCap at UW ITHS is supported by the National Center For Advancing Translational Sciences of the National Institutes of Health under Award Number UL1 TR002319. The content is solely the responsibility of the authors and does not necessarily represent the official views of the Society for Laboratory Automation and Screening or the National Institutes of Health. We would also like to thank Lochlan Hickok and Paul Miller for helping facilitate shipping and logistics; Molly W. Stephenson, Eden Anana, Jodie C. Tokihiro, Yixuan Zhou, Aryam Chhazal and Ben Mous for helping with device fabrication; and Ingrid Jeacopello for helping with sample preparation for biological experiments.

## Author contributions

EB, ABT and ST conceptualized the research UNL designed CandyCollect devices. UNL, AMM and WCT fabricated CandyCollect devices. DLH optimized and engineered the candy to act as a timer for collection. UNL, XS, DLH, WCT, AMM, MGT, GWH and TCC reviewed the literature and collaboratively designed experiments and the human subjects study. XS planned biological experiments. XS, WCT and AMM, conducted biological experiments and data collection. KNA, advised on work with human subjects and regulatory protocols. MGT, GWH and TCC, recruited participants, made a platform to screen participant eligibility, and packaged and shipped CandyCollects to research participants. ABT, ST, EB, UNL, XS, DLH, WCT, AMM, MGT, GWH and TCC interpreted the data. UNL, XS, DLH, WCT, AMM, MGT, GWH and TCC wrote sections of the manuscript. UNL, WCT and AMM made figures for the manuscript. ERW and GPD provided expertise on clinical relevance and sampling pathogens in saliva. UNL, XS, WCT, AMM, KNA, GPD, ERW, ST and ABT edited and revised the manuscript. ABT and ST supervised the research.

## Conflicts of interest

A.B.T. has ownership in Stacks to the Future, LLC and E.B. has ownership in Stacks to the Future, LLC, Tasso, Inc., and Salus Discovery, LLC. However, this research is not related to these companies.

