## Supplementary Information for "CandyCollect: At-home saliva sampling for capture of respiratory pathogens"

**Title: CandyCollect: At-home saliva sampling for respiratory pathogen capture**

**Figure S1:** Schematic of device dimensions.
**Figure S2:** Qualification of integrated density.
**Figure S3:** Analysis of *S. pyogenes* eluted from CandyCollect devices.
**Figure S4:** Agarose gel electrophoresis demonstrated high selectivity of the qPCR assay.
**Table S1:** Mass, diameter, thickness, and dissolving time of CandyCollects in Figure 4A.
**Table S2:** Mass, diameter, thickness, and dissolving time of CandyCollects in Figure 4B.

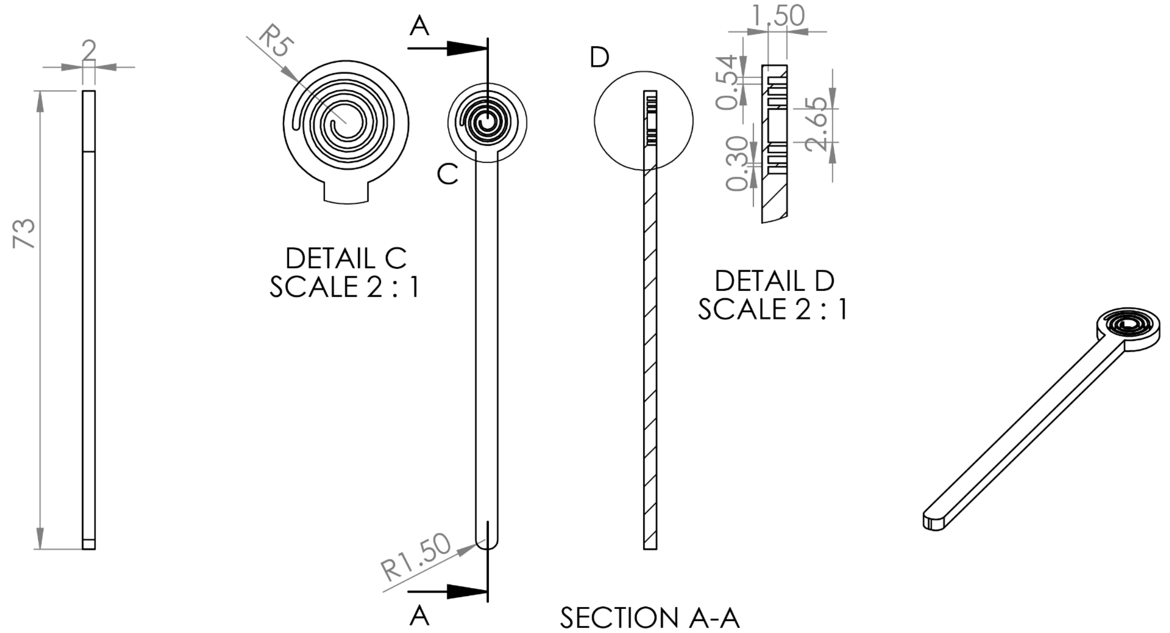


**Figure S1.** Schematic diagrams illustrating the dimensions, in mm, of the CandyCollect milled stick. CAD file also included.

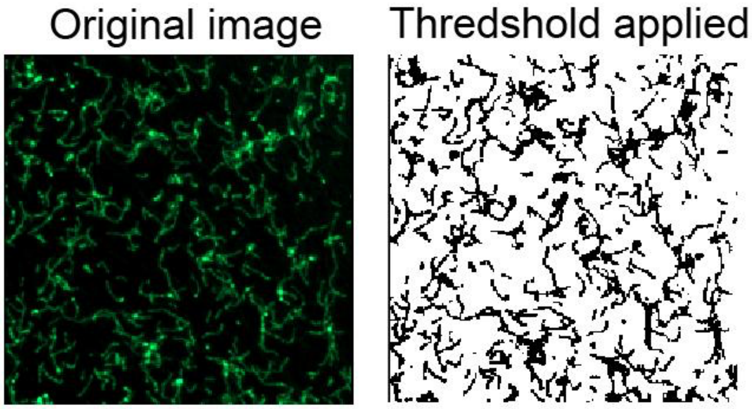


**Figure S2**. Qualification of integrated density. Image process before and after threshold applied in one given region of interest (ROI).

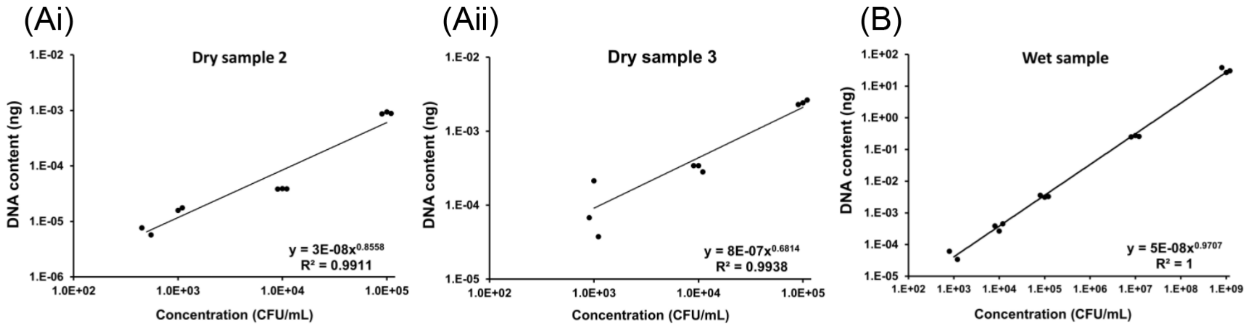


**Figure S3.** Analysis of *S. pyogenes* eluted from CandyCollect devices. (Ai-ii) Quantification of *S. pyogenes* from dry CandyCollect samples by qPCR. Shown here are two additional independent experiments performed in addition to the experiment shown in Figure 4C (note: the experiments in this figure were performed without enrichment of DNA, whereas DNA enrichment was performed in the experiment shown in Figure 4C). Each data point represents an individual CandyCollect. Three concentrations of *S. pyogenes* suspension $1x10^3$, $1x10^4$, $1x10^5$ CFU/mL were used. (B) Quantification of *S. pyogenes* from wet CandyCollect samples by qPCR. Each data point represents an individual CandyCollect. Five concentrations of *S. pyogenes* suspension $1x10^3$, $1x10^4$, $1x10^{5,}$ $1x10^{7,}$ $1x10^9$ CFU/mL were used. (Note: the procedures performed for the experiments shown in Figure S3A differed slightly from the procedure for Figure S3B; refer to the methods section for details.)

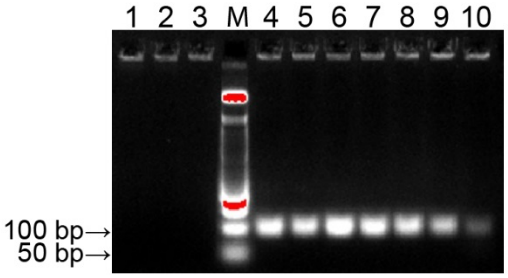


**Figure S4.** Agarose gel electrophoresis demonstrated high selectivity of the qPCR assay. qPCR was performed using the primers/probe targeting *spy1258*. The templates were 1:10 serial dilution of DNA (50 ng – 50 fg) from *S. pyogenes* and 50 ng of DNA from *S. mutans* and *S. aureus*. 3% agarose gel was used to separate DNA products from the qPCR reactions. A single PCR product with a size of about 100 bp, an expected amplicon size, was only shown in each of the DNA samples from *S. pyogenes* (lane 4-10). No PCR products were shown in DNA samples from *S. mutans* (lane 1), *S. aureus* (lane 2), and no template control (lane 3). Note: lane M is 50 bp DNA ladder.

**Table S1.** Mass, diameter, thickness, and dissolving time of CandyCollects in Figure 4A.

| Lollipop | Mass (g) | Diameter (mm) | Thickness (mm) | Time (min) |
|---|---|---|---|---|
| A1 | 0.04 | n/a* | n/a | 0.75 |
| A2 | 0.04 | n/a | n/a | 0.50 |
| A3 | 0.04 | n/a | n/a | 0.65 |
| B1 | 0.08 | n/a | n/a | 1.03 |
| B2 | 0.08 | n/a | n/a | 0.83 |
| B3 | 0.08 | n/a | n/a | 0.82 |
| C1 | 4.70 | 27.12 | 4.52 | 6.67 |
| C2 | 4.16 | 27.12 | 4.52 | 5.98 |
| C3 | 4.09 | 27.12 | 4.52 | 5.67 |
| D1 | 4.06 | 23.00 | 6.46 | 8.33 |
| D2 | 4.12 | 23.00 | 6.46 | 7.67 |
| D3 | 4.27 | 23.00 | 6.46 | 7.08 |
| E1 | 7.44 | 30.00 | 7.60 | 10.28 |
| E2 | 7.62 | 30.00 | 7.60 | 9.55 |
| E3 | 6.32 | 30.00 | 7.60 | 9.63 |
| F1 | 6.92 | 23.00 | 12.92 | 14.32 |
| F2 | 7.45 | 23.00 | 12.92 | 13.50 |
| F3 | 7.53 | 23.00 | 12.92 | 15.67 |

*Note: CandyCollects A1-B3 did not have measurable diameter and thickness because a mold was not used to apply the candy. Instead a small drop of isomalt was applied and allowed to form a thin layer on the stick. The diameter of the round area on the stick is 1 cm and the thickness is 2 mm.

**Table S2.** Mass, diameter, thickness, and dissolving time of CandyCollects in Figure 4B.

| Participant | Mass (g) | Diameter (mm) | Thickness (mm) | Time (min) |
|---|---|---|---|---|
| 1 | 0.96 | 16 | 4 | 2.08 |
| | 1.14 | 16 | 4 | 2.43 |
| | 1.07 | 16 | 4 | 2.40 |
| 2 | 0.93 | 16 | 4 | 3.42 |
| | 1.12 | 16 | 4 | 3.68 |
| | 0.97 | 16 | 4 | 1.97 |
| 3 | 1.02 | 16 | 4 | 4.77 |
| | 0.88 | 16 | 4 | 3.80 |
| | 1.07 | 16 | 4 | 4.43 |
| 4 | 1.07 | 16 | 4 | 5.28 |
| | 0.88 | 16 | 4 | 4.50 |
| | 1 | 16 | 4 | 4.22 |
| 5 | 1.06 | 16 | 4 | 2.83 |
| | 1.11 | 16 | 4 | 2.58 |
| | 1.09 | 16 | 4 | 2.45 |
| 6 | 0.98 | 16 | 4 | 2.72 |
| | 1.15 | 16 | 4 | 6.93 |
| | 0.97 | 16 | 4 | 4.62 |
| 7 | 1.07 | 16 | 4 | 2.35 |
| | 1 | 16 | 4 | 3.02 |
| | 1.09 | 16 | 4 | 3.13 |
| 8 | 1.12 | 16 | 4 | 2.50 |
| | 1 | 16 | 4 | 1.98 |
| | 0.96 | 16 | 4 | 2.08 |
| 9 | 1.13 | 16 | 4 | 2.55 |

| | | | | |
|---|---|---|---|---|
| | 1.03 | 16 | 4 | 2.27 |
| | 0.95 | 16 | 4 | 2.00 |
| 10 | 1.01 | 16 | 4 | 6.27 |
| | 0.83 | 16 | 4 | 5.98 |
| | 0.96 | 16 | 4 | 5.37 |
| 11 | 0.92 | 16 | 4 | 1.25 |
| | 1.04 | 16 | 4 | 1.33 |
| | 1.05 | 16 | 4 | 1.57 |
| 12 | 0.99 | 16 | 4 | 2.70 |
| | 1.02 | 16 | 4 | 2.53 |
| | 0.98 | 16 | 4 | 2.27 |
| 13 | 1.08 | 16 | 4 | 4.33 |
| | 0.95 | 16 | 4 | 4.07 |
| | 0.96 | 16 | 4 | 4.57 |
| 14 | 0.92 | 16 | 4 | 2.68 |
| | 1.01 | 16 | 4 | 2.60 |
| | 0.97 | 16 | 4 | 3.03 |
| 15 | 0.97 | 16 | 4 | 6.62 |
| | 1.02 | 16 | 4 | 7.70 |
| | 0.94 | 16 | 4 | 5.20 |
| 16 | 0.93 | 16 | 4 | 3.18 |
| | 0.95 | 16 | 4 | 3.13 |
| | 0.95 | 16 | 4 | 3.03 |
| 17 | 1.01 | 16 | 4 | 4.62 |
| | 1.04 | 16 | 4 | 4.47 |
| | 0.95 | 16 | 4 | 3.63 |
